# Gammaherpesvirus infection licenses age-associated B cells for pathogenicity in MS and EAE

**DOI:** 10.1101/2021.07.22.453263

**Authors:** Isobel C. Mouat, Jessica R. Allanach, Vina Fan, Anna M. Girard, Iryna Shanina, Galina Vorobeychik, Marc S. Horwitz

**Author notes:** Corresponding author: Marc S. Horwitz, Room 3551, Life Sciences Centre, 2350 Health Sciences Mall, University of British Columbia, Vancouver, B. C. Canada V6T 1Z3.

## Abstract

While age-associated B cells (ABCs) are known to expand and persist following viral infection and during autoimmunity, their interactions are yet to be studied together in these contexts. Epstein-Barr virus (EBV) infection has long been implicated in multiple sclerosis (MS), and it is not known whether ABCs could play a role in mediating viral contribution to autoimmunity. Here, we show that the circulating ABC population is expanded in people with MS and that EBV infection and MS status differentially impact the circulating ABC phenotype. We then directly compared ABCs during viral infection and autoimmunity using mouse models of EBV, gammaherpesvirus 68 (*γ*HV68), and MS, experimental autoimmune encephalomyelitis (EAE). We observed that splenic ABCs are expanded in a sex-biased manner during both latent virus infection and EAE, and each event drives the ABC population to opposing phenotypes. We have previously shown that latent *γ*HV68 infection exacerbates EAE and here we show that mice lacking ABCs fail to display *γ*HV68-enhanced disease. Collectively, these findings indicate that latent viral infection and central nervous system autoimmunity differentially impact the ABC population and suggests that viral infections such as EBV prime ABCs to contribute pathogenically in MS.

## Introduction

Age-associated B cells (ABCs) are a unique subset of immune B cells found in humans and mice. They have been identified and characterized independently in the contexts of female aging, viral infection, and autoimmunity(1–3). In life, these are overlapping and intertwined occurrences. In this study, we show that viral infection and autoimmunity differentially impact the phenotype of ABCs in humans and mice, and we identify ABCs as functional mediators of viral-enhanced autoimmunity.

ABCs, defined by high expression of CD11c and T-bet and low expression of CD21, are primarily found in the spleen and secrete antiviral or auto-antibodies, cytokines, and act as potent antigen-presenting cells(1,4,5). The number of ABCs increases with age, particularly in females(3,6–10). ABC numbers are also increased in people with autoimmune diseases, including systemic lupus erythematosus (SLE), rheumatoid arthritis (RA), individuals with multiple sclerosis (MS) over 60 years of age, and a subset of people with common variable immunodeficiency that display autoimmune complications(3,6,7,9–11). In a mouse model of SLE, knocking out T-bet specifically in B cells impairs the development of germinal centers, decreases autoantibody levels, and dampens kidney damage and mortality(12). ABCs have been shown to produce proinflammatory cytokines IFNγ and TNFα as well as regulatory cytokine IL-10 in autoimmune contexts(4,13). To our knowledge, ABCs have yet to be examined in the in vivo MS-model experimental autoimmune encephalomyelitis (EAE), though ABC differentiation depends on T-bet, a transcription factor that is essential for EAE(14). The functional capacities of ABCs and their contribution(s) to MS and EAE are not well understood. ABCs also expand in an array of viral infections including lymphocytic choriomeningitis virus (LCMV), murine cytomegalovirus (MCMV), gammaherpesvirus68 (*γ*HV68), vaccinia, human immunodeficiency virus (HIV), influenza, hepatitis C, and rhinovirus(15–22). ABCs have primarily been examined separately in the contexts of viral infection and autoimmunity. Given the distinct environments of viral infection and autoimmunity, it would be predicted that ABCs would take on distinct phenotypes and functional capacities. Recently, ABCs in people with HIV and SLE were shown to have similar but not completely overlapping transcriptional profiles(23). The phenotypic and functional characteristics of ABCs between disease states deserves further investigation.

Here, we do a side-by-side comparison of the ABC population in the contexts of both viral infection and autoimmunity in both humans and mice to ascertain the specific interactions driven by these two processes and their relative role in disease progression. We also examine the role of ABCs as mediators between gammaherpesvirus infection and MS/EAE. Epstein-Barr virus (EBV) is a highly prevalent virus, infecting over 90% of the adult human population(24). The association between EBV and autoimmunity was first demonstrated in 1979 when it was reported that peripheral blood lymphocytes from MS patients have an increased tendency to transform in vitro in response to EBV(25). Clinical findings support the association between EBV and MS: people with MS have higher titres of EBV-specific antibodies than healthy individuals(26) which correlate with disease activity(27), and T cells in people with MS display aberrant responses to EBV(28). Further work has demonstrated that EBV’s association with autoimmune disease extends to SLE and RA(29–34). The precise mechanism of EBV’s contribution to MS is controversial – EBV-infection of autoreactive B cells, molecular mimicry, bystander activation of autoreactive cells, EBV-infected B cell invasion of the central nervous system (CNS) and activation of Human Endogenous Retrovirus-W, and EBV-induced cytokine response have all been suggested(35,36).

Our group has previously demonstrated that infection with latent *γ*HV68, a murine model of EBV(37), exacerbates MOG_35-55_ EAE in C57BL/6(J) mice(38). Mice latently infected with *γ*HV68 display an earlier and more severe EAE clinical course without viral reactivation or CNS infection. Latently *γ*HV68-infected mice have a heightened peripheral cytotoxic T cell response and develop CNS lesions composed of CD4^+^ and CD8^+^ T cells, with a proportion of infiltrating CD8^+^ T cells specific for *γ*HV68. Disease enhancement is dependent on the development of latency and is specific to *γ*HV68; other viral infections including LCMV and MCMV do not lead to EAE enhancement(39). This *γ*HV68-EAE model is a valuable model of the relationship between EBV and MS and here, we demonstrate that the specific induction of ABCs is required for the latent *γ*HV68-enhancement of EAE.

In this study, we directly compare the ABC population in the contexts of both latent viral infection and autoimmunity in humans and mice and distinguish the role of ABCs between different models of CNS autoimmunity, EAE and latent *γ*HV68 enhancement of EAE. We propose that alteration of the ABC population due to gammaherpesvirus infection is a mechanism by which viral infection contributes to the development and progression of autoimmune disease.

## Results

### EBV and MS status alter the circulating ABC population

We first compared the ABC population frequency and phenotype in individuals with and without a history of EBV infection and MS. Peripheral blood mononuclear cells (PBMCs) were collected from EBV^-^ and EBV^+^ healthy women and people with MS, all of whom are EBV^+^ women, and the ABC proportion and phenotype were examined by flow cytometry. The people diagnosed with MS were not exposed to immunomodulatory treatments and had disease durations ranging from 4 months to 7 years (Table 1). All donors were female and median ages between groups were comparable: EBV^-^ healthy donors (25 years), EBV^+^ healthy donors (26.5 years), and EBV^+^ people with MS (31 years) (Table 1). EBV and CMV serostatus was determined by ELISA.

**Table 1:** Donor characteristics.

| Age at donation (years) | Country of birth and age of immigration to Canada (if applicable) | Self-identified ethnicity (boxes checked) | Age (years) at MS symptom onset | EDSS score at time of donation | Disease duration at time of donation | EBV status | CMV status |
| --- | --- | --- | --- | --- | --- | --- | --- |
| 25 | Country: Canada<br>Age: N/A | Canadian | 23 | 2.0 | 2 years 8 months | VCA+, EBNA-1+ | - |
| 24 | Country: United Kingdom<br>Age: 5 years | Middle Eastern | 24 | 2.5 | 4 months | VCA+, EBNA-1+ | + |
| 31 | Country: Afghanistan<br>Age: 20 years | Middle Eastern | 31 | 2.5 | 6 months | VCA+, EBNA-1- | + |
| 32 | Country: Canada<br>Age: N/A | Canadian | 26 | 2.0 | 5 years | VCA+, EBNA-1+ | - |
| 39 | Country: Canada<br>Age: N/A | Canadian, English, Scottish, Irish | 31 | 2.0 | 7 years | VCA+, EBNA-1+ | - |
| 25 | Country: Canada<br>Age: N/A | Canadian, English | N/A | N/A | N/A | VCA-EBVNA1- | - |
| 26 | Country: Canada<br>Age: N/A | Canadian | N/A | N/A | N/A | VCA-EBVNA1- | - |
| 18 | Country: Canada<br>Age: N/A | Canadian, Chinese | N/A | N/A | N/A | VCA-EBVNA1- | + |
| 39 | Country: Canada<br>Age: N/A | Canadian | N/A | N/A | N/A | VCA+ EBNA1+ | - |
| 23 | Country: Canada<br>Age: N/A | Canadian, Scottish | N/A | N/A | N/A | VCA+ EBNA1+ | + |
| 25 | Country: Canada<br>Age: N/A | Canadian, Scottish, Swedish, First Nations | N/A | N/A | N/A | VCA+ EBNA1- | + |
| 28 | Country: Canada<br>Age: N/A | Canadian, Dutch | N/A | N/A | N/A | VCA+ EBNA1+ | - |

For this study, ABCs were defined as CD19^+^CD20^+^CD11c^+^CD21^-^ (Figure 1A). Gene analysis has shown CD19^+^CD21^-^ B cells display high levels of CD11c and express T-bet(7,40). Further, CD11c and CD21 have been used by other groups to define ABCs in people(3,6). Only extracellular staining was performed to preserve cell numbers due to low prevalence of ABCs.

**Figure 1:**
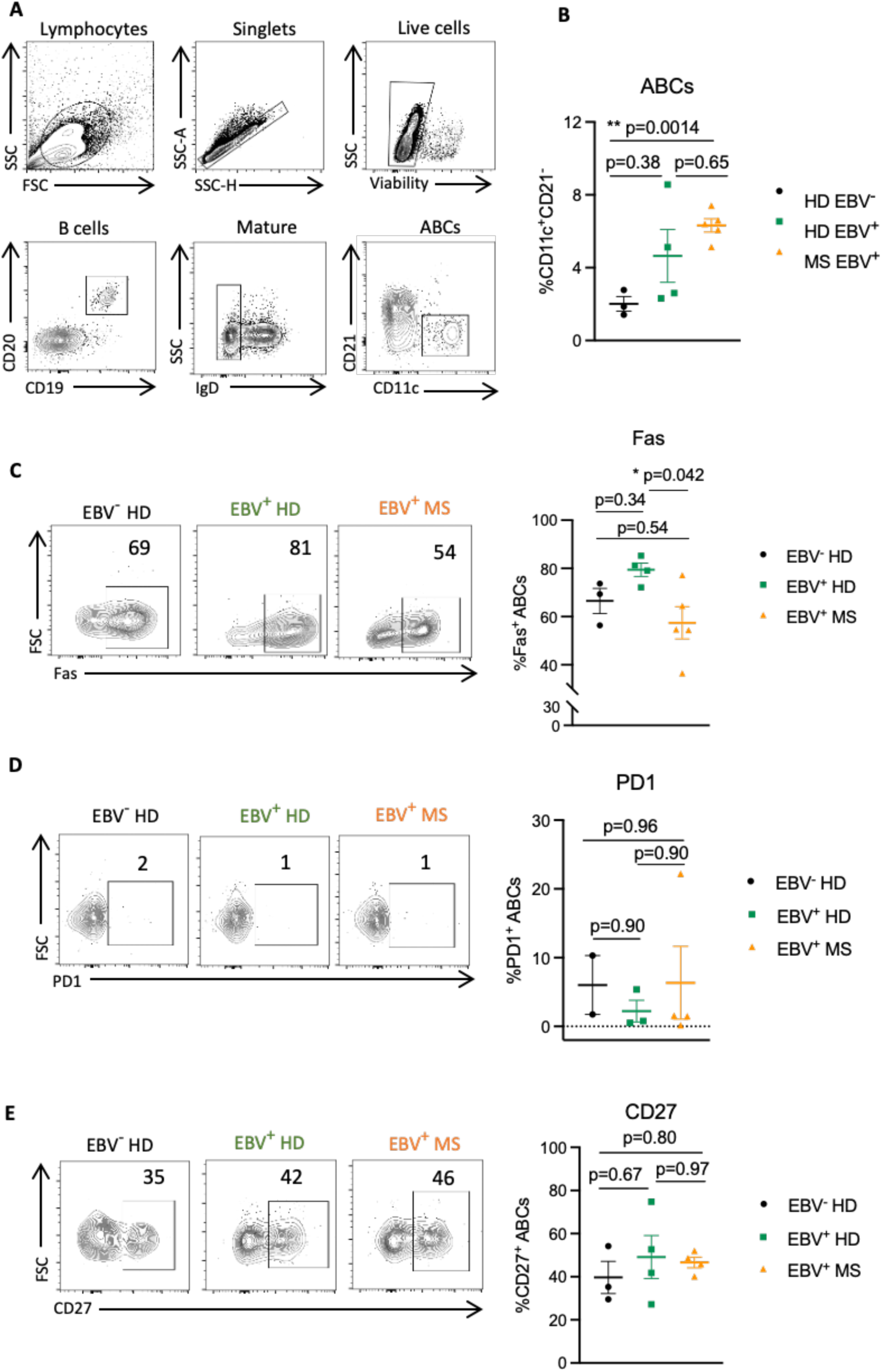
The peripheral ABC phenotype is impacted by EBV infection and MS status. PBMCs were collected from three EBV seronegative healthy females (EBV^-^ HD, black filled circles), four EBV seropositive healthy females (EBV^+^ HD, green filled squares), and five females with relapse-remitting MS that are all EBV seropositive (EBV^+^ MS, orange filled triangles). PBMCs were processed for flow cytometry immediately following isolation. (**A**) Representative gating strategy for ABCs (CD19^+^CD20^+^CD11c^+^CD21^-^) from PBMCs. (**B**) Proportion ABCs (CD11c^+^CD21^-^) among mature B cells (CD19^+^CD20^+^IgD^-^). Representative samples and percent of ABCs (CD19^+^CD20^+^CD11c^+^CD21^-^) positive for (**C**) Fas (**D**) PD1, and (**E**) CD27. Data for CD27 and PD1 stained samples are fewer due to later addition of these markers following initial findings. Data presented as mean ± SEM, analyzed by one-way ANOVA with multiple comparisons, ** p<0.01, * p<0.05.

We found that people with MS have increased proportions of circulating ABCs compared to EBV seronegative healthy people (Figure 1B). EBV seropositive healthy individuals show a nonsignificant trend towards increased ABC proportions compared to EBV seronegative healthy people but reduced compared to MS (Figure 1B). Among the donor groups, we do not observe differences in the proportion of total circulating B cells (Supplementary Figure 1A). These findings indicate that both EBV infection and MS might expand the proportion of circulating ABCs. In addition to changes to ABC proportions, we also observe differences in ABC phenotype between groups.

We examined the impact of EBV infection and MS on the ABC phenotype, because the impact of autoimmunity versus viral infection on the ABC phenotype has not been thoroughly studied. For these purposes, our preliminary analysis focused on the proportion of ABCs expressing Fas, PD1, and CD27. ABCs have previously been shown to express substantially more Fas than non-ABC B cells(3,40) and ABCs in people with common variable immune deficiency express more Fas than ABCs in healthy donors(9). Another inhibitory receptor, PD1, has also been shown to be upregulated on ABCs compared to other B cell subsets both in healthy individuals and those with SLE(10,16). The role of CD27, a marker of memory, on ABCs is not well understood – ABCs have been reported to express high CD27(3), though some groups define ABCs as CD27 negative(16,40). It is not known how infection versus autoimmunity impact the expression of Fas, PD1, and CD27 on ABCs. We observed that slightly over half of ABCs express Fas, and EBV infection led to a nonsignificant increase in the proportion of ABCs positive for Fas compared to EBV seronegative healthy individuals (Figure 1C). Among people who are EBV seropositive, MS led to a significant decrease in the proportion of ABCs expressing Fas compared to healthy donors (Figure 1C). In contrast to previous reports(10,16), we found that a low proportion of ABCs express PD1 and the proportion of ABCs that express PD1 is not different across EBV seronegative or EBV seropositive healthy individuals or people with MS (Figure 1D). Finally, just under half of ABCs express CD27, and EBV and MS status did not impact the proportion of ABCs that express CD27 (Figure 1E).

We observed that CMV status did not significantly correlate with the ABC phenotype in healthy individuals nor those with MS (Supplemental Figure 1), indicating that CMV, another herpesvirus linked to MS, may be less influential of this B cell population compared to EBV. To further investigate how ABCs differ between the contexts of viral infection and MS, and whether ABCs can act as mediators of the viral contribution to disease, we investigated in vivo models of infection and disease.

### *γ*HV68 infection and EAE induction increase ABCs

To determine whether ABCs are differentially impacted by gammaherpesvirus infection and an in vivo model of MS, we examined the proportion of ABCs following latent *γ*HV68 infection and EAE induction. We examined the relative proportions of ABCs in C57BL/6(J) mice mock-infected (naïve), infected i.p. with 10^4^ PFU *γ*HV68 for 35 days to establish latency (*γ*HV68), induced with EAE (MOG_35-55_ with pertussis toxin on day 0 and 2 post-induction) for 12-14 days (EAE), or infected with *γ*HV68 for 35 days, then induced for MOG_35-55_ EAE for 12-14 days (*γ*HV68-EAE). We collected spleens, brains, and spinal cords, processed for, and performed flow cytometry to examine ABCs (Supplemental Figure 2A). We asked whether the abundance of ABCs in the spleen changes with *γ*HV68 infection, EAE induction, or both, and whether ABCs are found at the site of disease, the CNS, during EAE.

We found that ABCs expand in the spleen in a sex-biased manner during *γ*HV68 infection and EAE. While *γ*HV68 infection increases the relative proportion of ABCs among mature B cells in the spleen in both male and female mice, induction of EAE increases the proportion of ABCs only in females (Figure 2A-B). During *γ*HV68-EAE, the proportion of ABCs are increased compared to naïve mice in both females and males (Figure 2A-B). The proportion of ABCs is increased in females as compared to males in *γ*HV68, EAE, and *γ*HV68-EAE mice, though there is no sex bias observed in naïve mice (Figure 2C-F). We did not observe a difference in T-bet expression in splenic ABCs between groups (Supplemental Figure 2B). Due to the observed sex bias, we focused our analysis on female mice.

**Figure 2:**
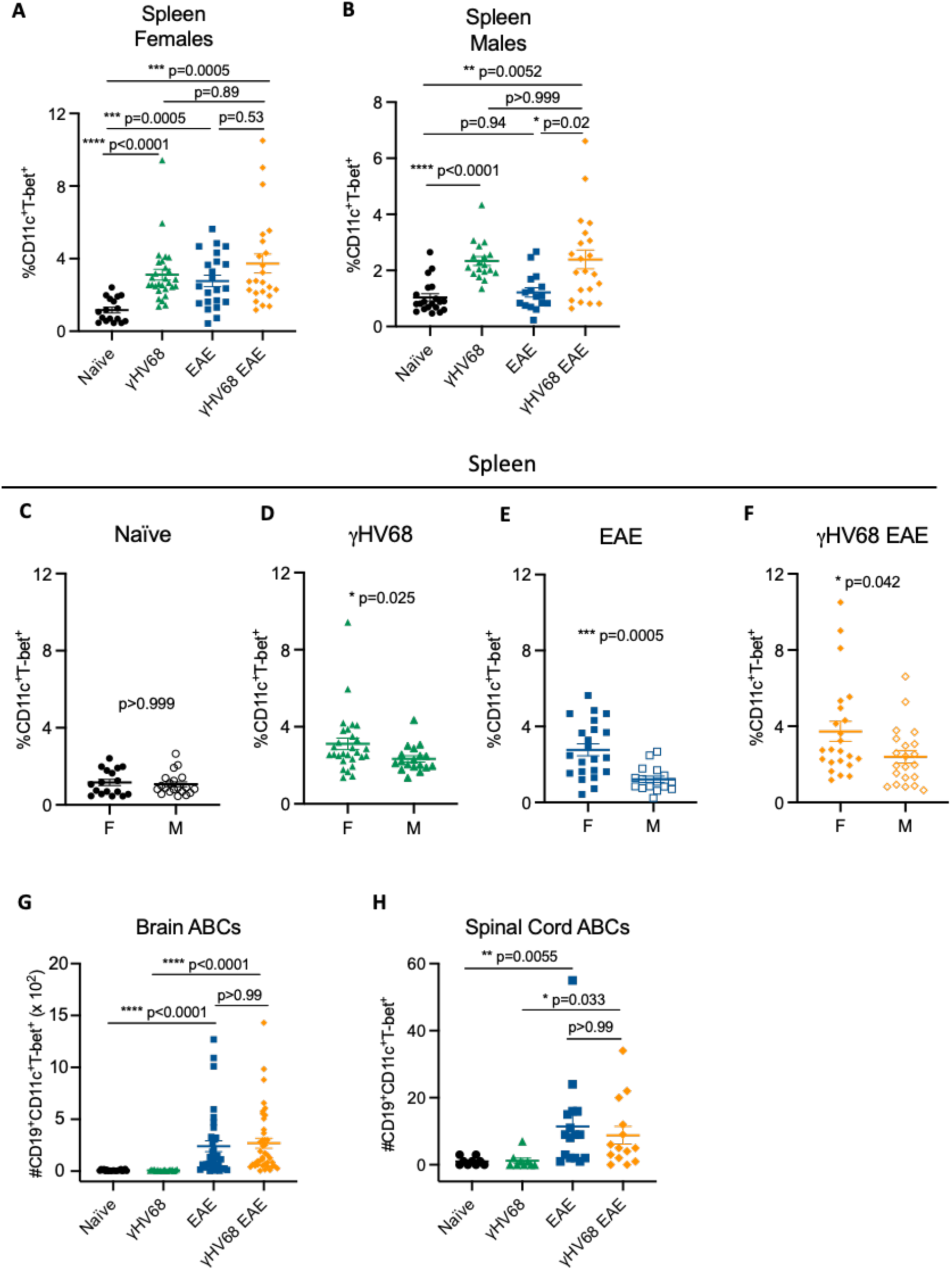
ABCs are expanded in a sex-biased manner during γHV68 infection and EAE. C57BL/6(J) mice infected with γHV68 for 35 days to establish latency (γHV68, green triangles) or mock-infected with blank medium (naïve, black circles). In some mice, MOG_35-55_ EAE was induced 35 days after mock infection (EAE, blue squares) or initial infection (γHV68 EAE, orange diamonds). At 35 days p.i. or 13 days post EAE induction, spleen, brain, and spinal cord collected and processed for flow cytometry. (**A-B**) Proportion of ABCs (CD11c^+^T-bet^+^) of mature B cells (CD19^+^IgD^-^) in the spleen of females and males. (**C-F**) Proportion of ABCs (CD11c^+^T-bet^+^) of mature B cells (CD19^+^IgD^-^) in the spleen of females (F) and males (M). Data repeated from panels A-B in panels C-F for better visualization and comparison of sex differences. (**G-H**) Number of ABCs in the brain(**G**) and spinal cord(**H**). (**A-F**) Data pooled across 9 experiments, n=16-29 per group. (**G-H**) Data pooled across 5 experiments, n=9-41 per group. Data presented as mean ± SEM, analyzed by one-way ANOVA (**A, B**), Mann-Whitney (**C-F**), or Kruskal-Wallis (**G, H**) test, ****p<0.0001, *** p<0.001; ** p<0.01, * p<0.05.

We asked whether *γ*HV68-infection or EAE leads changes ABC numbers in the CNS, the site of disease during EAE. Not surprisingly, under control conditions where CNS infiltration does not occur, specifically, *γ*HV68 infection alone and without EAE induction, did not result in increased numbers of ABCs in the brain or spinal cord (Figure 2G-H). However, EAE induction increased the number of ABCs in the brain and spinal cord in both uninfected and *γ*HV68-infected mice (Figure 2G-H). Unlike the spleen, we did not observe a sex difference in the number of ABCs within the CNS (Supplemental Figure 2C-D). We found that ABCs constitute a lower proportion of total lymphocytes in the brain and spinal cord compared to the spleen (Supplemental Figure 2E), indicating that they are likely not preferentially homing to the CNS during disease. The precise localization of the CNS-infiltrating ABCs is not known, and whether they localize in the meninges, parenchyma, or associate with lesions or ectopic lymphoid structures, could give insight into their role within the CNS during disease.

These findings demonstrate that ABCs display a sex bias in the contexts of viral infection and autoimmunity and support previous findings that ABCs are largely resident in the spleen. To further investigate the role(s) of ABCs in viral infection and autoimmunity, we next examined the phenotype of ABCs in these different contexts.

### *γ*HV68 infection and EAE differentially impact ABC phenotype

Although ABCs are implicated in both viral infection and autoimmunity, to our knowledge the ABC phenotype has not been directly examined in these two instances. To compare how *γ*HV68 infection and EAE induction influence the ABC phenotype, we used flow cytometry to examine extracellular and intracellular antigens on splenic ABCs to compare the phenotype of ABCs specifically in EAE with and without prior *γ*HV68 infection. We found that *γ*HV68 infection and EAE drive differential ABC phenotypes that are distinct from each other and the ABC phenotype of naïve mice. Specifically, *γ*HV68 infection led to an increase in the proportion of ABCs expressing IFN*γ* and TNF*α*, and a decrease in ABCs expressing IL-17A (Figure 3A, Supplemental Figure 3A-C). In contrast, EAE results in an increased proportion of ABCs expressing IL-17A and IL-10, and no change in IFN*γ* or TNF*α*, compared to ABCs in naïve mice (Figure 3A, Supplemental Figure 3A-C).

**Figure 3:**
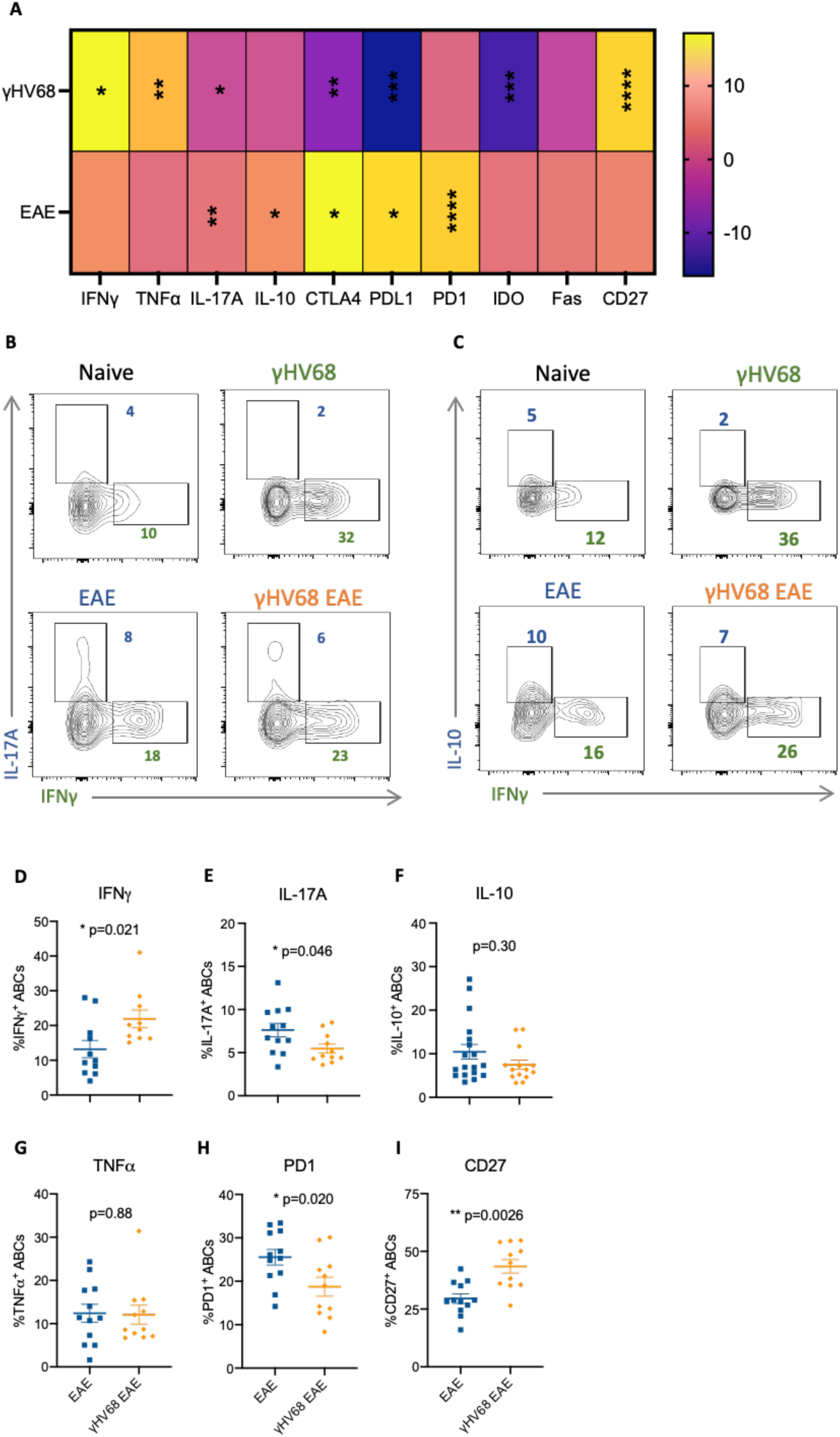
γHV68 infection and EAE drive differential ABC phenotypes. C57BL/6(J) mice infected with γHV68 for 35 days (γHV68) or mock-infected (naïve). In some mice, MOG_35-55_ EAE was induced 35 days after mock infection (EAE, blue squares) or initial infection (γHV68 EAE, orange diamonds). At 35 days p.i. or 13 days post EAE induction, spleen collected and processed for flow cytometry. (**A**) Heatmap – changes in frequency of ABC population expressing given markers compared relative to naïve controls. (**B-C**) Representative flow plots. Previously gated on CD19^+^CD11c^+^T-bet^+^ ABCs in the spleen. Concatenated files. (**D-I**) Percent of splenic ABCs (CD19^+^CD11c^+^T-bet^+^) that are positive for (**D**) IFNγ, (**E**) IL-17A, (**F**) IL-10, (**G**) TNF*α*, (**H**) PD1, (**I**) CD27. Data pooled across 2-3 experiments, n=9-18 per group. (**A**) Analyzed by one-way ANOVA with multiple comparisons, compared to naïve, (**D-I**) data presented as mean ± SEM, analyzed by Mann-Whitney test, ****p<0.0001, *** p<0.001, ** p<0.01, * p<0.05.

Further, *γ*HV68 infection results in significantly fewer ABCs expressing the inhibitory receptors CTLA4, PDL1, and IDO, while EAE increases the proportion of ABCs expressing CTLA4, PD1, and PDL1 (Figure 3A, Supplemental Figure 3E-H). The proportion of ABC expressing Fas appeared unchanged by *γ*HV68 infection or EAE (Figure 3A, Supplemental Figure 3I). *γ*HV68 infection increased the proportion of ABCs expressing the memory marker CD27, while EAE did not (Figure 3A, Supplemental Figure 3J).

We asked how *γ*HV68 infection prior to EAE induction impacts the phenotype of ABCs. We observed that the ABCs in *γ*HV68-EAE mice display an intermediate phenotype between those of *γ*HV68 and EAE mice. In particular, the proportion of ABCs in *γ*HV68-EAE mice positive for cytokines IL-17A, IFN*γ*, and IL-10 is intermediate between those of *γ*HV68 and EAE mice (Figure 3B-C, Supplemental Figure 3A, C, D). In addition, an intermediate amount of the ABCs in *γ*HV68-EAE mice expressed inhibitory receptors CTLA4, PDL1, PD1, and IDO between those in *γ*HV68 and EAE (Supplemental Figure 3E-H). This intermediate phenotype displayed by ABCs in *γ*HV68-EAE mice appeared to be more pathogenic than the ABC phenotype in uninfected EAE mice. We found that ABCs in *γ*HV68-EAE mice express more IFN*γ* and less IL-17A than ABCs in EAE mice (Figure 3D-E, Supplemental Figure 3A, C), indicating that ABCs in *γ*HV68-EAE mice display a more Th-1 associated phenotype than those in EAE mice. We detected no difference in the proportion of ABCs expressing IL-10 or TNF*α* between EAE and *γ*HV68-EAE mice, though ABCs in *γ*HV68-EAE express less PD1 and more CD27 (Figure 3F-I, Supplemental Figure 3G, J). As we have previously shown that *γ*HV68-exacerbation of EAE is associated with a robust Th-1 response, in both the periphery and CNS, the presence of the Th-1 like ABC phenotype suggested that *γ*HV68-primed ABCs could contribute to the exacerbation of EAE disease.

### ABC knockout mice demonstrate that ABCs are protective in EAE and pathogenic in *γ*HV68-EAE

Next, we sought to determine whether ABCs contribute to the EAE or *γ*HV68-EAE clinical course. Previously, our lab has shown that latent *γ*HV68 infection prior to disease induction results in an exacerbated clinical course, with a Th-1 skewed immune response and increased CD8 T cell infiltration to the CNS(38).

To investigate the role of ABCs in disease we examined mice with a B cell specific T-bet deletion(12) that lack ABCs. We demonstrate a significant loss of ABCs in *Tbx21^fl/fl^Cd19^cre/+^* (KO) compared to *Tbx21^fl/fl^Cd19^+/+^* (Ctrl) littermate controls (Figure 4A). Female Ctrl and KO mice were mock-infected or infected with *γ*HV68 for 35 days and induced for MOG_35-55_ EAE. Mice were monitored daily post EAE-induction, blinded to genotype, and measured on a 5-point scale as previously described(38).

**Figure 4:**
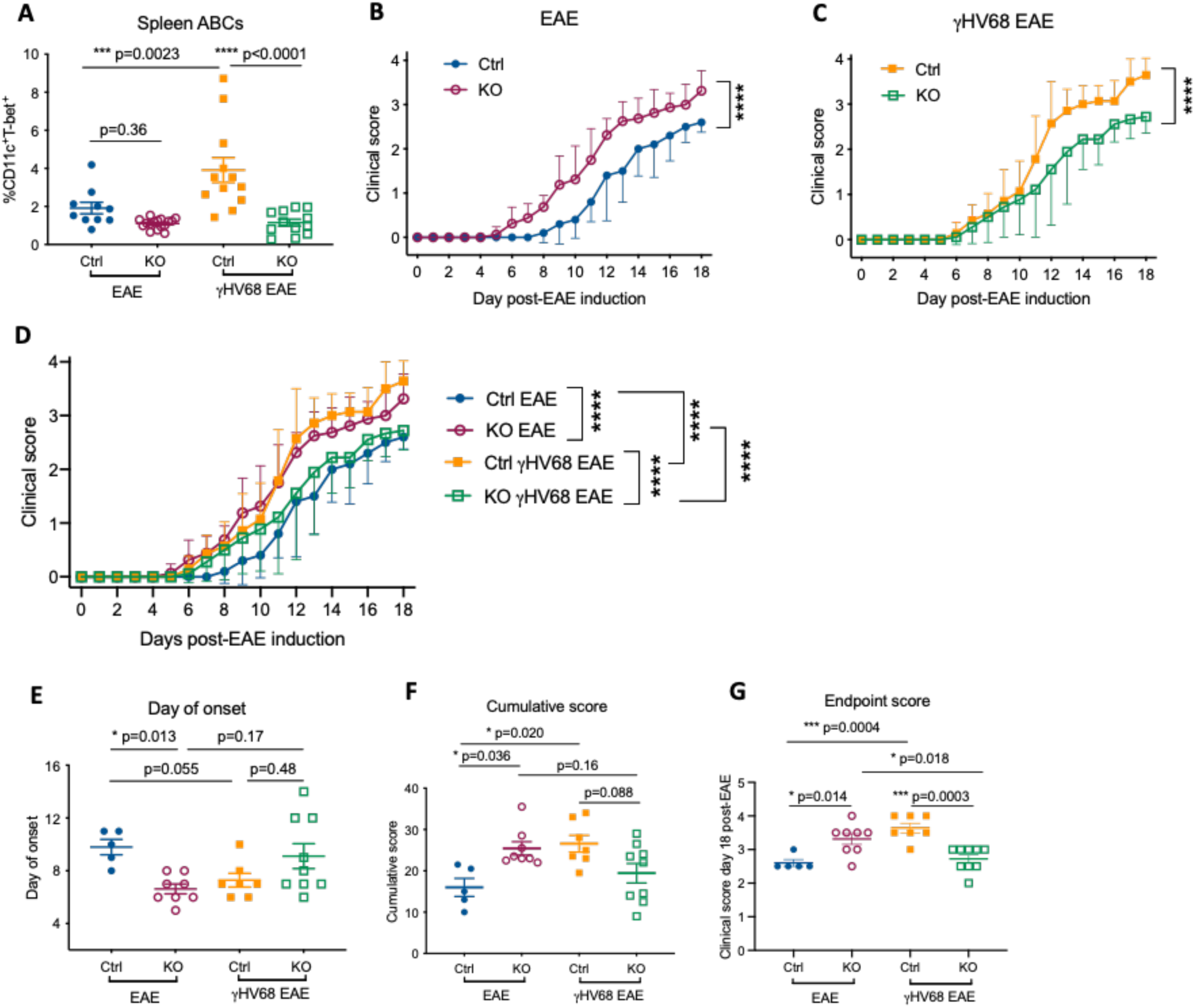
EAE disease course is altered in ABC KO mice. Female *Tbx21^fl/fl^Cd19^cre/+^* (KO) and *Tbx21^fl/fl^Cd19^+/+^* (Ctrl) mice were infected with γHV68 or mock-infected for 35 days and then induced for MOG_35-55_ EAE and clinically monitored for 18 days. (**A**) Percent ABCs (CD11c^+^T-bet^+^) of mature B cells (CD19^+^IgD^-^) in the spleen. (**B-D**) EAE clinical scores. Data in panel D is repeated from panels B-C. Data are pooled across three experiments, n=5-9 per group, representative of five independent experiments. (**E**) Day of clinical symptom onset. (**F**) Cumulative score (**G**) endpoint score at day 18 post-EAE induction. Data analyzed by (**A, E-G**) one-way ANOVA, (**B-C**) two-way ANOVA, (**D**) two-way ANOVA with multiple comparisons. Data presented as mean ± SEM (**A, E-G**) or mean ± SD (**B-D**), ****p<0.0001, *** p<0.001, ** p<0.01, * p<0.05.

Strikingly, we observed that mice lacking ABCs had the opposite effect on the clinical course of EAE and *γ*HV68-EAE. Specifically, during EAE ABC KO mice developed more severe disease than Ctrl mice, while in *γ*HV68-EAE, the disease course was less severe than Ctrl mice (Figure 4B-D). During EAE, KO mice displayed an earlier day of onset, and higher cumulative and endpoint scores than Ctrl mice (Figure 4E-G). In contrast, *γ*HV68-EAE KO mice displayed a lower endpoint score (Figure 4G).

These results indicate that ABCs contribute protectively in EAE and pathogenically in *γ*HV68-EAE and these results align with our prior observations that ABCs display a more pathogenic phenotype in *γ*HV68-EAE, compared to EAE. In addition, while in Ctrl mice *γ*HV68-EAE resulted in an exacerbated disease course compared to EAE alone, ABC KO mice did not display enhanced CNS disease thereby demonstrating that ABCs are required for *γ*HV68-enhancement of EAE.

### Knocking out ABCs impacts the T cell population but not autoantibodies

ABCs are major producers of IgG2a/c(18,41) and, in a model of SLE, knocking out ABCs results in a significant decrease in anti-chromatin IgG2a/c titres(12). To ask whether autoantibodies were mechanistically responsible for the impact in disease observed here, we analyzed the sera from these mice. Surprisingly, we did not observe differences in anti-MOG IgG2a/c titres between Ctrl and KO mice during EAE and *γ*HV68-EAE (Figure 5A-B), indicating that, in this context, ABCs are contributing to disease via a mechanism other than autoantibody production.

**Figure 5:**
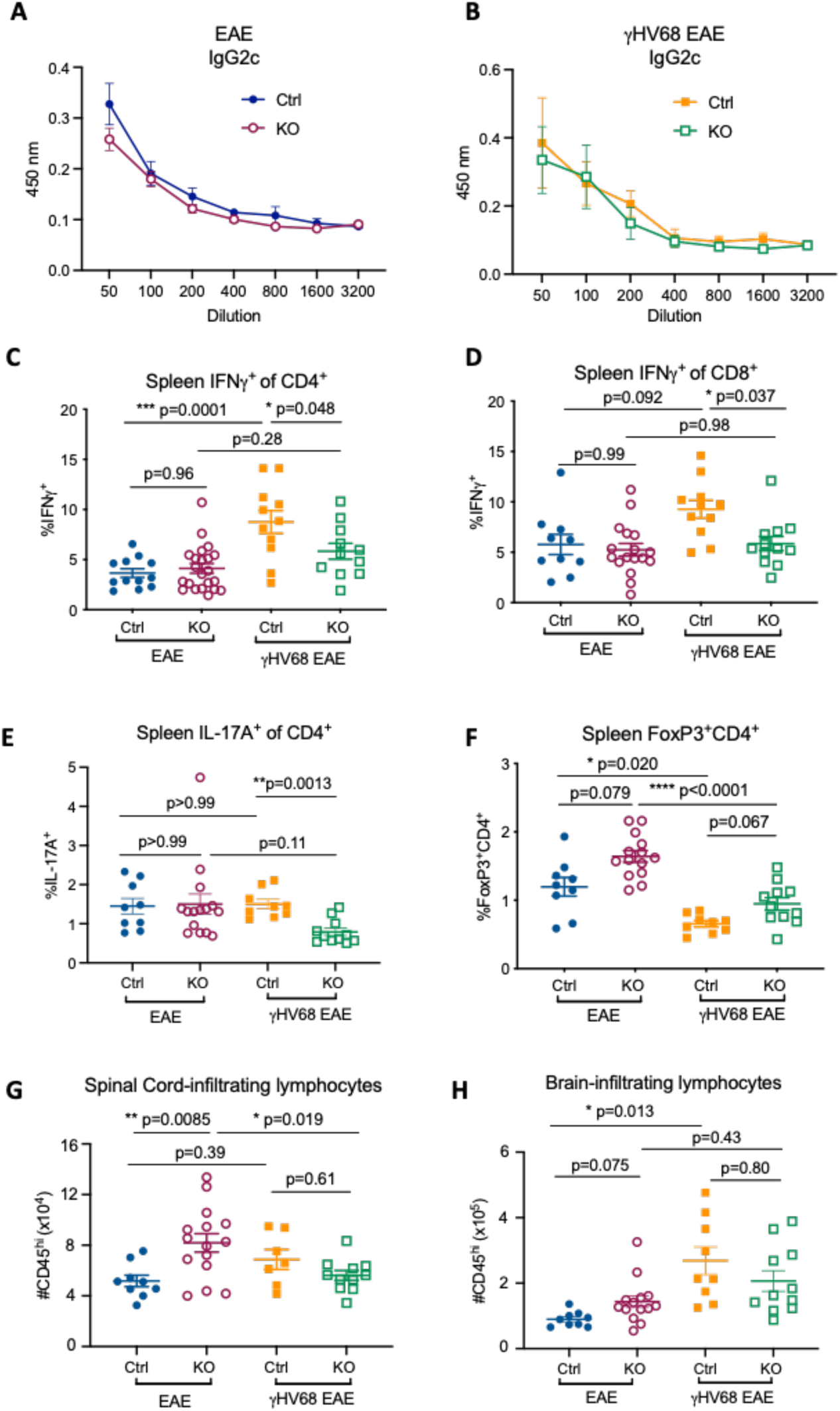
Immune profile, but not autoantibodies, is altered in ABC KO mice. Female *Tbx21^fl/fl^Cd19^cre/+^* (KO) and *Tbx21^fl/fl^Cd19^+/+^* (Ctrl) mice were infected with γHV68 or mock-infected for 35 days and then induced for MOG_35-55_ EAE. At day 13-18 post-EAE induction blood (serum), spleen, brain, and spinal cord collected. Spleen, brain, and spinal cord processed for flow cytometry. (**A-B**) Optical density (O.D., y-axis) reflecting titers (x-axis; dilution of serum) of IgG2c anti-MOG_35-55_ antibodies in Ctrl and KO mice, separated by EAE (**A**) and γHV68 EAE (**B**) mice, n=3-6 mice per group. (**C-D**) Percent IFNγ^+^ of CD3^+^CD4^+^ (**C**) and CD3^+^CD8^+^ (**D**) cells in the spleen. (**E**) Percent of CD3^+^CD4^+^ cells in the spleen expressing IL-17A. (**F**) Percent of live cells in the spleen that are CD4^+^FoxP3^+^. (**G-H**) Number of CD45^hi^ cells in the spinal cord (**G**) and brain (**H**). Data presented as mean ± SEM. (**C-H**) Analyzed by one way ANOVA with multiple comparisons, ****p<0.0001, *** p<0.001, ** p<0.01, * p<0.05.

It is known that ABCs are capable of presenting antigen(5) and we and others see that ABCs are predominantly spleen-resident(17). Therefore, we speculated that ABCs could impact T cell polarization in the spleen and infiltration of the CNS. In the spleen there is no difference in the proportion of cytotoxic or helper T cells between Ctrl and KO mice during EAE or *γ*HV68-EAE (Supplemental Figure 4A-B). However, the skewing of the peripheral T cells is altered in KO mice during *γ*HV68-EAE. During *γ*HV68-EAE, KO mice did not display the increase in IFN*γ*-expressing CD4 or CD8 T cells in the spleen that is seen in Ctrl mice (Figure 5C-D). Additionally, significantly fewer helper T cells express IL-17A in KO mice during *γ*HV68-EAE compared to Ctrl mice(Figure 5E). The decrease in proportion of IFN*γ*^+^ and IL-17A^+^ T cells in KO mice during *γ*HV68-EAE could explain the dampened disease course in these mice. In addition, there were proportionately fewer CD4 T cells expressing IL-10 in KO mice during *γ*HV68-EAE (Supplemental Figure 4C). The decrease in proportion of T cells expressing IFN*γ* and IL-17A and increase in IL-10 likely indicates that, without ABCs, the peripheral immune response is shifting towards a tolerogenic state. KO mice in both EAE and *γ*HV68-EAE displayed a trend of increased regulatory T cells in the spleen, indicating that ABCs may dampen the regulatory T cell population (Figure 5F).

In contrast to *γ*HV68-EAE mice, there were no changes in splenic T cell skewing during EAE between Ctrl and KO mice (Figure 5C-D). In EAE mice, the proportion of cytotoxic and helper T cells, and the proportion of CD4 T cells expressing IFN*γ*, IL-17A, and IL-10 did not differ between Ctrl and KO mice (Figure 5C-E, Supplemental Figure 4A-C). These results indicate that ABCs impact T cell skewing in the spleen during *γ*HV68-EAE, though not EAE.

In EAE mice, although we did not observe major differences in the periphery between Ctrl and KO mice, there were significant differences in CNS lymphocyte infiltration. KO mice induced for EAE displayed increased numbers of lymphocytes infiltrating the spinal cord, where disease predominates, compared to Ctrl (Figure 5G). This indicates that rather than impacting T cell skewing in the periphery, ABCs in EAE mice could possibly be influencing CNS lymphocyte homing or blood-brain barrier integrity, yet there was no significant difference in CNS-infiltrating lymphocytes in *γ*HV68-EAE (Figure 5G-H). Further, there were no significant differences in the number of cytotoxic, helper, or regulatory T cells infiltrating the CNS between Ctrl and KO mice during EAE or *γ*HV68-EAE (Supplemental Figure 4D-I). While eliminating some cell types, these results beg further analysis to understanding the cell populations that are infiltrating the CNS in KO mice during EAE.

Prior work had shown that specifically deleting T-bet from B cells lead to an inability of mice to control LCMV chronic infection(18,41). To test the impact of *γ*HV68 replication, we found an increased quantity of *γ*HV68 in the spleen of ABC KO mice compared to Ctrl during *γ*HV68-EAE (Supplemental Figure 4J). The ability of ABCs to control *γ*HV68 load and the clinical course of disease warrants further investigation.

Together, these findings indicate that ABCs influence T cell skewing in the periphery and lymphocyte infiltration of the CNS during disease. The precise mechanisms by which ABCs impact T cell skewing and lymphocyte migration to and infiltration of the CNS warrants further investigation.

## Discussion

In this report, we have shown that that age-associated B cell population is differentially impacted by viral infection and autoimmunity, in both mice and humans. We have sought to directly compare the ABC population in these two contexts and examine ABCs in conditions of combined latent viral infection and autoimmunity, which is relevant to MS patients. In addition, we examined the functional contribution of ABCs and identified them as a mediator between latent gammaherpesvirus and MS disease.

Together, these findings indicate that ABCs are differentially impacted by viral infection and autoimmunity, in both mice and people. Additionally, we found that ABCs in *γ*HV68-EAE mice take on a phenotype with characteristics of both the viral infection and disease, demonstrating that ABCs are a heterogeneous population, at least phenotypically, particularly when challenged by multiple antigens. These findings have implications for studying the human ABC population that is undoubtedly impacted by an array of pathogens and contexts.

A surprising finding of this study was the female sex bias of ABCs in mice infected with *γ*HV68 and/or induced for EAE. This finding is relevant to the question of if ABCs play a role in the sex-bias observed in various autoimmune diseases, including MS and SLE, as has previously been suggested(42). Interestingly, MOG_35-55_ EAE does not display a sex bias in C57BL/6 mice(43,44), so if and how the increased numbers of ABCs in female EAE mice impact disease in this model remains unknown.

We observed that the ABC phenotype is differentially impacted by viral infection and autoimmunity in mice and humans, though the phenotype observed in mice and humans does not always align. For instance, we observe that the proportion of ABCs that express Fas is decreased in people with MS compared to healthy EBV^+^ donors, while in mice we observe an increase in *γ*HV68-EAE compared with *γ*HV68. This disconnect between the mouse and human ABC phenotype may be due to intrinsic differences between the immune systems in mice and humans or the lack of previous antigen exposure in the mice. Additionally, it must be considered whether circulating ABCs are representative of those in the spleen. We and others observed that ABCs are predominantly spleen-resident in mice(17), and further investigation should examine ABCs in the spleen versus circulation, especially because most human samples examined are blood.

We also found that ABCs are heterogenous in terms of phenotype and functional capacity depending on context and can contribute to disease in a protective or pathogenic manner. ABCs likely comprise several subsets that differ in terms of functional capacity. In the future, it will be important to determine disease-associated subsets; a portion of ABCs could contribute to disease, and others act in a regulatory/protective manner. An outstanding question is whether the ABCs population is effectively depleted by Rituximab, and if the ABC population is affected by other immunomodulatory treatments.

Interestingly, it appears that ABCs contribute to some models of autoimmunity in a protective manner, and others pathogenically. We observed that ABCs contribute to EAE in a protective manner, as knocking them out results in an exacerbation of disease, while in a model of SLE ABCs appear to be contributing pathogenically(12). We have previously found that knocking out ABCs has no impact on the clinical course of collagen-induced arthritis, though mice without ABCs do not show the *γ*HV68-exacerbation of arthritis that is observed in control mice(45). The finding that ABCs are required for the *γ*HV68 exacerbation of both CIA and EAE brings up the question of whether ABCs are a conserved mechanism by which gammaherpesvirus contributes to both arthritis and MS.

Critical to understanding this discrepancy in ABCs functioning protectively versus pathogenically in different models of autoimmunity is determining the mechanisms by which ABCs impact disease. Precisely how ABCs contribute to autoimmunity disease is unclear, and they may be contributing to different diseases through different mechanisms. It is intriguing that in a model of SLE, knocking out ABCs results in substantial changes to autoantibodies, yet we see no change in autoantibodies in models of EAE and CIA. Instead, we show here that during EAE and *γ*HV68-EAE, knocking out ABCs results in altered peripheral T cell responses and lymphocyte infiltration of the CNS. In this context, ABCs may be functioning more as antigen-presenting cells rather than antibody-secreting cells. ABCs are known to be potent antigen presenting cells and B cells contribute to EAE through their antigen presentation functions(46).

It remains unknown how ABCs are impacting disease in the CNS during *γ*HV68-EAE, as we do not observe differences in lymphocyte number or phenotype in KO mice. Other groups have found that immune cell infiltration numbers and histopathological markers of CNS damage do not reliably correlate with EAE severity(47,48). The increased *γ*HV68 load in KO mice might indicate viral reactivation. Our lab has previously shown that only latent *γ*HV68, not lytic, leads to exacerbation of EAE(38). Whether the virus is reactivating, and, if so, if the presence of the lytic virus is modulating disease, deserves further examination.

In summary, we have shown that 1) EBV status and MS diagnosis impact the proportion of circulating ABCs and their phenotype, 2) ABCs expand in the spleen in a sex-biased manner during *γ*HV68 infection and EAE, 3) ABCs display distinct phenotype during *γ*HV68 infection and EAE, and 4) knocking out ABCs results in exacerbation of EAE disease however loss of *γ*HV68-enhancement of EAE. We hope that these findings can contribute to better understanding of the relationship between EBV and MS.

## Methods

### Donor recruitment and enrolment

Five individuals with relapsing MS were recruited at the Burnaby Hospital MS Clinic (Fraser Health) under the supervision of Dr. Galina Vorobeychik. Seven unaffected donors were recruited at the Life Sciences Center (University of British Columbia). All donors were female, 19 – 39 years of age (mean age 28 ± 6.4 years) and provided written informed consent prior to enrolment in the study. Donors with a definite RRMS diagnosis, according to Poser or 2010 McDonald criteria, and disease duration of less than 10 years, were confirmed as treatment naïve prior to donation (no previous use of any disease modifying therapies during lifetime). MS donors underwent a neurological exam the day of blood donation to assess Expanded Disability Status Scale (EDSS). Individuals with a progressive MS diagnosis or EDSS >4, that were male, pregnant, outside the designated age range, or undergoing treatment were excluded from the present study.

### Donor blood collection and processing

Blood samples were obtained by venipuncture and assigned an alphanumeric identifier. Whole blood was processed for peripheral blood mononuclear cell (PBMC) isolation by Lymphoprep (StemCell) gradient separation, according to manufacturer’s instructions, within an hour of collection in K2-EDTA coated vacutainer tubes (BD). Freshly isolated PBMC were counted and stained for flow cytometric analysis (no prior freezing or other preservation steps used). Serum was obtained by centrifugation of untreated blood samples and frozen at -80°C for subsequent analysis.

### Donor serology

Antibodies to EBV and CMV were detected using enzyme-linked immunosorbent assays (ELISA) to determine donor history of infection. Nunc Maxisorp 96-well microtiter ELISA plates (Thermo Fisher Scientific) were coated overnight at 4°C with 1 μg/well of the peptide of interest in carbonate buffer. For EBV EBNA-1 and CMV, epitope peptides 1 and 2 were mixed together prior to well coating. The following day, the plates were washed with PBS and wash buffer (PBS + 0.05% Tween-20), followed by a two-hour blocking step at room temperature (RT) using wash buffer + 3% bovine serum albumin. Plates were washed again, then incubated with serum samples diluted in blocking buffer. Serum diluted 1:1000 was used to determine seropositivity to EBV and CMV antigens. After a two-hour RT incubation with serum, the plates were washed, then incubated with HRP-labelled goat anti-human IgG antibody (Thermo Fisher Scientific) diluted 1:3000 in blocking buffer or HRP-labelled goat anti-human IgM (Thermo Fisher Scientific) diluted 1:4000 at 37°C for 1 hour. The plates were then washed with PBS prior to the addition of 100 μL/well TMB substrate (BD Biosciences). Fifteen minutes after the addition of substrate, 100 μL stop solution (2 M sulfuric acid) was added to each well. The plates were read at 450 nm on a VarioSkan Plate Reader (Thermo Fisher Scientific) within 10 minutes of adding stop solution.

| Antigen | Peptide sequence |
| --- | --- |
| EBV VCA p18 | ASAGTGALASSAPSTAVAQSATPSVSSSISSLRAATSGATAAAVDTGS<br>GGGGQPHDTAPRGARKKQ |
| EBV EBNA-1 | Epitope 1: RSPSSQSSSSGSPRRPPPGRRPFFHPVG<br>Epitope 2: DYFEYHQEGGPDGEPDVPPGAIEQGPADDPGEGPSTGPRG |
| CMV | Epitope 1: CETDDLDEEDTSIYLSPPPVPVQVVAKRLRPDTPRT<br>Epitope 2: KSGTGPQPGSAGMGGAKTPSDAVQNILQKIEKIKNTEE |

### Mice

C57BL/6(J) mice were purchased from the Jackson Laboratory*. Tbx21^fl/fl^Cd19^cre/+^* mice were generated by crossing *Tbx21^fl/fl^Cd19^cre/+^* and *Tbx21^fl/fl^Cd19^+/+^* mice. *Tbx21^fl/fl^* and *Cd19^cre/+^* mice were provided by Dr. Pippa Marrack(12). All animals were bred and maintained in the specific-pathogen free animal facility at the University of British Columbia. All animal work was performed in accordance with the regulation of the Canadian Council for Animal Care.

### *γ*HV68 infection

*γ*HV68 (WUMS strain, purchased from ATCC) was propagated on Baby Hamster Kidney (BHK, ATCC) cells and quantified by plaque assay. Mice were infected with *γ*HV68 or mock-infected with Minimum Essential Media (MEM, Gibco). For infection, virus was diluted in MEM and maintained sterile and on ice until infection. 6- to 10-week-old mice were infected intraperitoneally (i.p.) with 10^4^ plaque-forming units (PFU) *γ*HV68. No clinical symptoms were observed from viral infection.

### Induction and evaluation of EAE

EAE was induced 35 days post-*γ*HV68 infection by injecting 100 μl of emulsified Complete Freund’s Adjuvant (DIFCO) containing 200 μg of MOG_35-55_ (GenScript) and 400 μg of desiccated Mycobacterium tuberculosis H37ra (DIFCO) subcutaneously. Mice also received two doses of 200 ng of pertussis toxin (List Biologicals) i.p. at the time of immunization and 48 hours post-induction. Mice were assessed daily (blinded) on a scale from 0 to 5: 0, no clinical symptoms, 0.5, partially limp tail; 1, paralyzed tail; 2, loss of coordination; 2.5, one hind limb paralyzed; 3, both hind limbs paralyzed; 3.5, both hind limbs paralyzed accompanied by weakness in the forelimbs; 4, forelimbs paralyzed (humane endpoint); 5, moribund or dead.

### Tissue harvesting

Mice were euthanized by cardiac puncture while anesthetized with isoflurane and immediately perfused with 20cc PBS to allow for brain and spinal cord harvesting without blood contamination. Spleen, brain, and spinal cord were extracted, placed into PBS, and temporarily kept on ice until processing. Spleens and spinal cords were mashed through a 70 μm cell strainer and a single cell suspension prepared for each sample. Brains were mashed through a 70 μm cell strainer twice and a single-cell suspension was prepared for each sample. Splenocytes were incubated in Ammonium-Chloride-Potassium (ACK) lysing buffer for 10 minutes on ice to lyse red blood cells and remaining cells were kept on ice until further use. Immune cells were isolated from brains and spinal cords using a 30% Percoll gradient (GE Healthcare) and were resuspended in Flow Cytometry Staining Buffer (FACS buffer, PBS with 2% newborn calf serum, Sigma-Aldrich) until staining.

### Flow cytometric analysis of cell-type specific surface antigens and intracellular cytokines

To evaluate cytokine production by various cell types, 4 million splenocytes cells were stimulated ex vivo for 3 hours at 5% CO2 at 37°C in Minimum Essential Media (Gibco) containing 10% fetal bovine serum (FBS, Sigma-Aldrich), 1 μl/ml GolgiPlug (BD Biosciences), 10 ng/ml PMA (Sigma-Aldrich) and 500 ng/ml ionomycin (Thermo Fisher Scientific). Stimulated cells were then washed prior to staining. For each spleen sample, 4 million cells were stained in 2 wells of a 96-well plate, with 2 million cells per well. All collected brain and spinal cord cells were resuspended in FACS buffer and stained in a single well. Blood donor PBMC samples were stained at 1 million cells per well using the following method without ex vivo stimulation. Prior to staining, samples were incubated for 30 minutes at 4°C with 2 μl/ml Fixable Viability Dye eFluor506 (Thermo Fisher Scientific) while in PBS. Cells were then incubated with a rat anti-mouse CD16/32 (Fc block, BD Biosciences) antibody, or Human BD Fc Block (BD Biosciences) for PBMC samples, for 10 minutes at RT. Fluorochrome labeled antibodies against cell surface antigens were then added to the cells for 30 minutes covered from light at 4°C. After washing, cells were resuspended in Fix/Perm buffer (Thermo Fisher Scientific) for 30 minutes-12 hours covered from light minutes at 4°C, washed twice with Perm buffer, and incubated 40 minutes with antibodies against intracellular antigens in Perm buffer. Cells were then washed and resuspended in FACS buffer with 2 mM EDTA. Staining with antibodies from Thermo Fisher Scientific, Biolegend, and BD Biosciences. Samples collected on Attune NxT Flow Cytometer (Thermo Fisher Scientific). Analyzed with FlowJo software version 10 (FlowJo LLC). Full-minus-one controls used for gating.

| Anti-human antibodies |  |  |  |
| --- | --- | --- | --- |
| Antigen | Fluorophore | Clone | Company |
| CD19 | APC-eFluor 780 | HIB19 | Thermo Fisher Scientific |
| IgD | FitC | IA6-2 | BD Bioscience |
| CD11c | AlexaFluor 700 | B-ly6 | BD Bioscience |
| CD21 | PerCpCy5.5 | Bu32 | Biolegend |
| CD20 | Pacific Blue | 2H7 | Biolegend |
| CD27 | Pe-Cy7 | LG.3A1 | Biolegend |
| Fas | PE Texas Red | DX2 | BD Bioscience |
| PD1 | PE | MIH4 | Thermo Fisher Scientific |

| Anti-mouse antibodies |  |  |  |
| --- | --- | --- | --- |
| Antigen | Fluorophore | Clone | Company |
| CD19 | PE or FitC | ID3 | Thermo Fisher Scientific |
| CD11c | APC | N418 | Biolegend |
| CD11c | Alexafluor700 | N418 | Thermo Fisher Scientific |
| T-bet | PerCpCy5.5 or PE | 4B10 | Thermo Fisher Scientific |
| IgD | Pe-Cy7 | 11-26c | Thermo Fisher Scientific |
| IFNγ | APC | XMG1.2 | Thermo Fisher Scientific |
| TNFα | Pe-Cy7 | MP6-Xt22 | Biolegend |
| IL-10 | Pacific Blue | JES5-16E3 | Biolegend |
| IL-17A | PE-Dazzle594 | TCII-18H10.1 | Biolegend |
| CTLA4 | PE-Dazzle594 | UC10-4B9 | Biolegend |
| PDL1 | SB436 | MIH5 | Thermo Fisher Scientific |
| PD1 | SB436 | J43 | Thermo Fisher Scientific |
| IDO | PerCpeFluor 710 | mIDO-48 | Thermo Fisher Scientific |
| Fas | FitC | SA367H8 | Biolegend |
| CD27 | Pe-Cy7 | LG.7F9 | Biolegend |
| CD45 | PerCpCy5.5 | 30-F11 | Thermo Fisher Scientific |
| CD4 | AlexaFluor 700 | RM4-5 | Thermo Fisher Scientific |
| CD8 | APC-eFlour 780 | 53-6.7 | Thermo Fisher Scientific |
| FoxP3 | FitC | FJK-16S | Thermo Fisher Scientific |
| CD3e | eFluor 450 | 500A2 | Thermo Fisher Scientific |

### Anti-MOG antibody ELISA

The sera were isolated by centrifugation 2000 × g for 10 min, aliquoted, and stored for up to 6 months at -80°C prior to running the ELISA. Anti-MOG_35-55_ antibodies were quantified by standard indirect ELISA. Briefly, ELISA plates (NUNC, Thermo Fisher Scientific) were coated with 1 μg MOG_35-55_ resuspended in coating buffer (0.05M carbonate buffer (pH 9.6)) overnight at 4°C. They were then washed 4x with wash buffer (PBS, 0.05% Tween-20), blocked with 5% newborn calf serum (NBCS, Sigma-Aldrich) for 1 hour at 37°C, incubated with serial dilutions (1:50 to 1:3200) of test sera diluted in blocking buffer for 2 hours at 37°C, and washed 4x wash buffer. Bound (anti-MOG_35-55_) antibody was incubated with HRP-conjugated goat anti-mouse IgG2c (Thermo Fisher) diluted 1:500 in blocking buffer, for 1 hour at 37°C, washed 4x with wash buffer, and detected by TMB substrate addition (BD Biosciences). The plates were read at 450 nm on a VarioSkan Plate Reader (Thermo Fisher Scientific) within 10 minutes of adding stop solution.

### *γ*HV68 quantitation

Quantification of *γ*HV68 load was done as previously described (49). gDNA was extracted from 4 × 10^6^ splenocytes with PureLink Genomic DNA mini kit (Thermo Fisher Scientific), according to the manufacturer’s instructions, and stored at -20°C. For quantitative qPCR, 150 ng DNA per reaction was amplified in duplicate using primers and probes specific to *γ*HV68 *Orf50* and mouse *Ptger2* (see table) and 2× QuantiNova Probe Mastermix (Qiagen). Standard curves were obtained by serial dilutions of *Orf50* and *Ptger2* gBlocks (ORF50: 2×10^6^ – 2×10^1^; PTGER2: 5×10^7^-5×10^2^). Run on the Bio-Rad CFX96 Touch™ Real Time PCR Detection system.

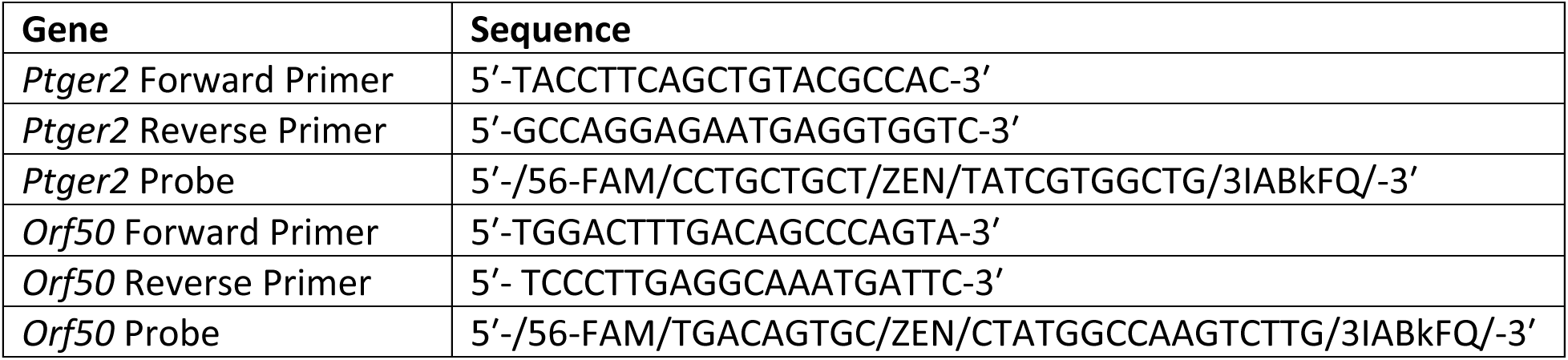

### Statistical analyses

Data and statistical analyses were performed using GraphPad Prism software 8.4.2 (GraphPad Software Inc.). Results are presented as mean ± SEM. Number of mice per group (n), statistical significance (p-value), and the number of experimental replicated as indicated in figure legend. Statistical analyses included: two-way ANOVA with Geisser-Greenhouse’s correction, Mann-Whitney test, or one-way ANOVA. P-values indicated by asterisks as follows: ****p<0.0001, *** p<0.001, ** p<0.01, * p<0.05.

### Study approval

Animal work was approved by the Animal Care Committee (ACC) of the University of British Columbia (Protocols A17- 0105, A17-0184). The human blood donation protocol (H16-02338) was approved by the University of British Columbia Clinical Research Ethics Board and by the Fraser Health Authority, General Neurology/Multiple Sclerosis.

## Author contributions

ICM and MSH conceptualized project and designed experiments. ICM performed mouse experiments, analyzed data, and wrote the manuscript. JRA processed human donor samples and contributed to the manuscript. VF serotyped blood donors. AMG assisted with experiments and developed MOG ELISA protocol. IS genotyped mice and assisted with experiments. GV provided human donor samples.

## Acknowledgements

We are grateful to Dr. Philippa Marrack for providing *Tbx21^fl/fl^* and *Cd19^cre/+^* mice. This research was supported in part by the MSSC, CIHR, UBC, and JDRF. We express our gratitude to all the blood donors who participated in this study.

## Supplement

**Supplemental Figure 1:**
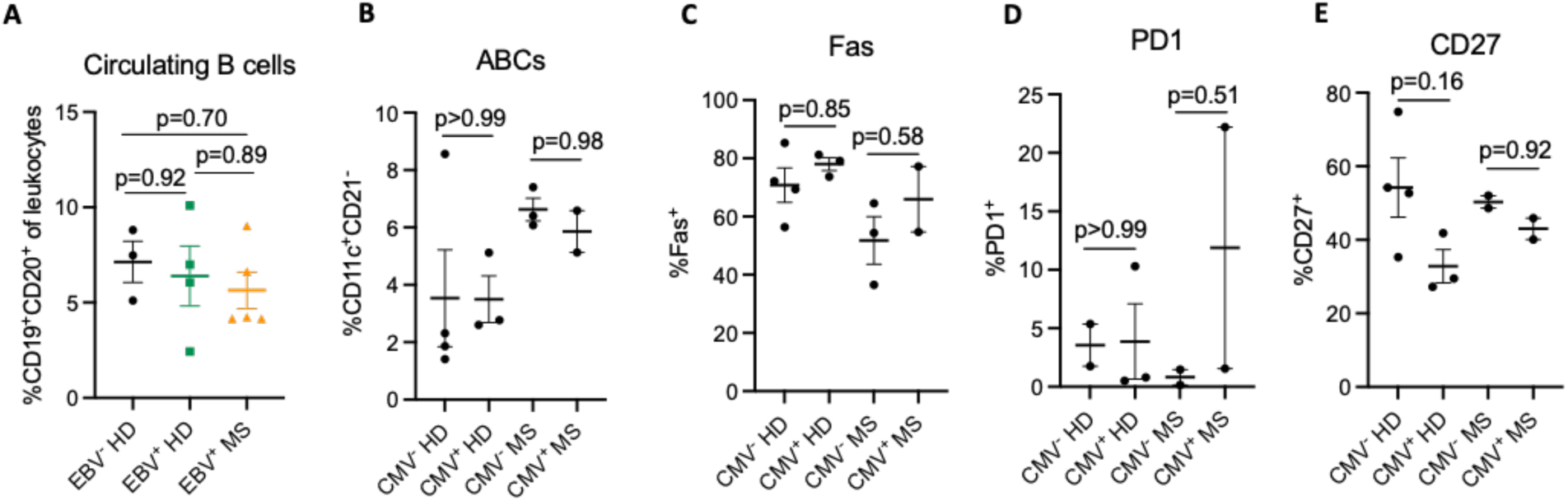
ABC phenotype is not impacted by CMV status. Same donors as Figure 1, here classified by CMV, rather than EBV status. PBMCs were collected from four CMV seronegative healthy females (CMV^-^ HD), three CMV seropositive healthy females (CMV^+^ HD), three CMV seronegative females with relapse-remitting MS (CMV^-^ MS), and two CMV seropositive females with relapse-remitting MS (CMV^+^ MS). PBMCs were processed for flow cytometry immediately following isolation. (**A**) Proportion of B cells (CD19^+^CD20^+^) of leukocytes, separated by EBV and MS status. (**B**) Proportion ABCs (CD11c^+^CD21^-^) among mature B cells (CD19^+^CD20^+^IgD^-^), separated by CMV status. (**C-E**) Percent of ABCs (CD19^+^ CD20^+^CD11c^+^CD21^-^) positive for (**C**) Fas, (**D**) PD1, and (**E**) CD27. Data presented as mean ± SEM, analyzed by one-way ANOVA with multiple comparisons.

**Supplemental Figure 2:**
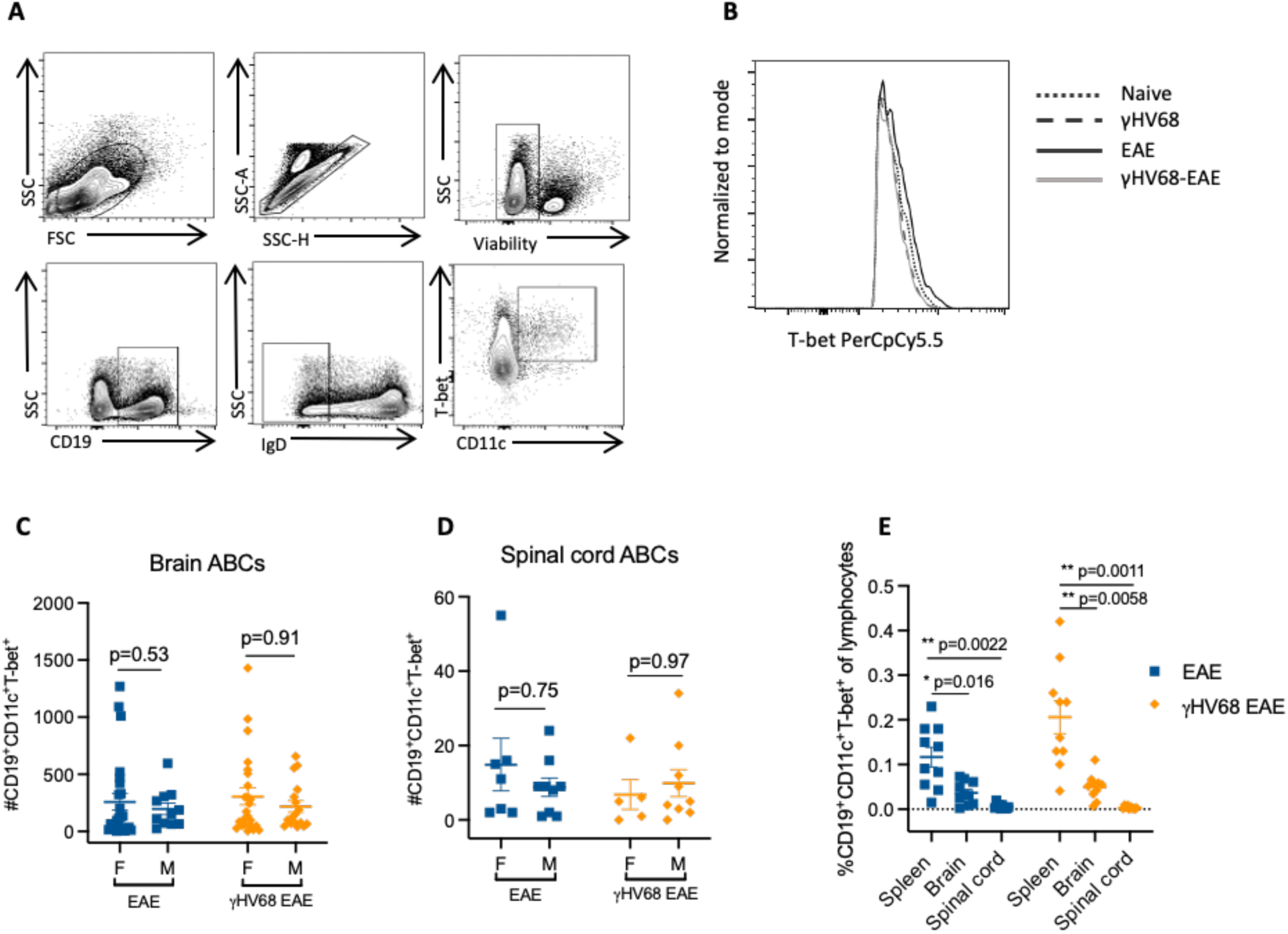
(**A**) Representative gating of ABCs: lymphocytes, singlets, live cells, CD19^+^ B cells, IgD^-^ mature B cells, CD11c^+^T-bet^+^ ABCs. (**B**) Representative expression of T-bet on ABCs (CD19^+^CD11c^+^T-bet^+^) in the spleen from naïve, latently γHV68-infected (35 days p.i.), EAE (13-15 days post-induction), and γHV68-EAE (13-15 days post-induction). (**C-D**) Number of ABCs (CD19^+^CD11c^+^T-bet^+^) in the brain (**C**) or spinal cord (**D**) of mice induced for EAE (13-15 days post-induction) and γHV68-EAE (13-15 days post-induction), as determined by flow cytometry. (**C-D**) Data pooled across 5 experiments, n=5-25 per group. (**E**) Percent of ABCs of live leukocytes in the spleen, brain, and spinal cord. Data presented as mean ± SEM, analyzed by one-way ANOVA test, *** p<0.001; ** p<0.01, * p<0.05.

**Supplemental Figure 3:**
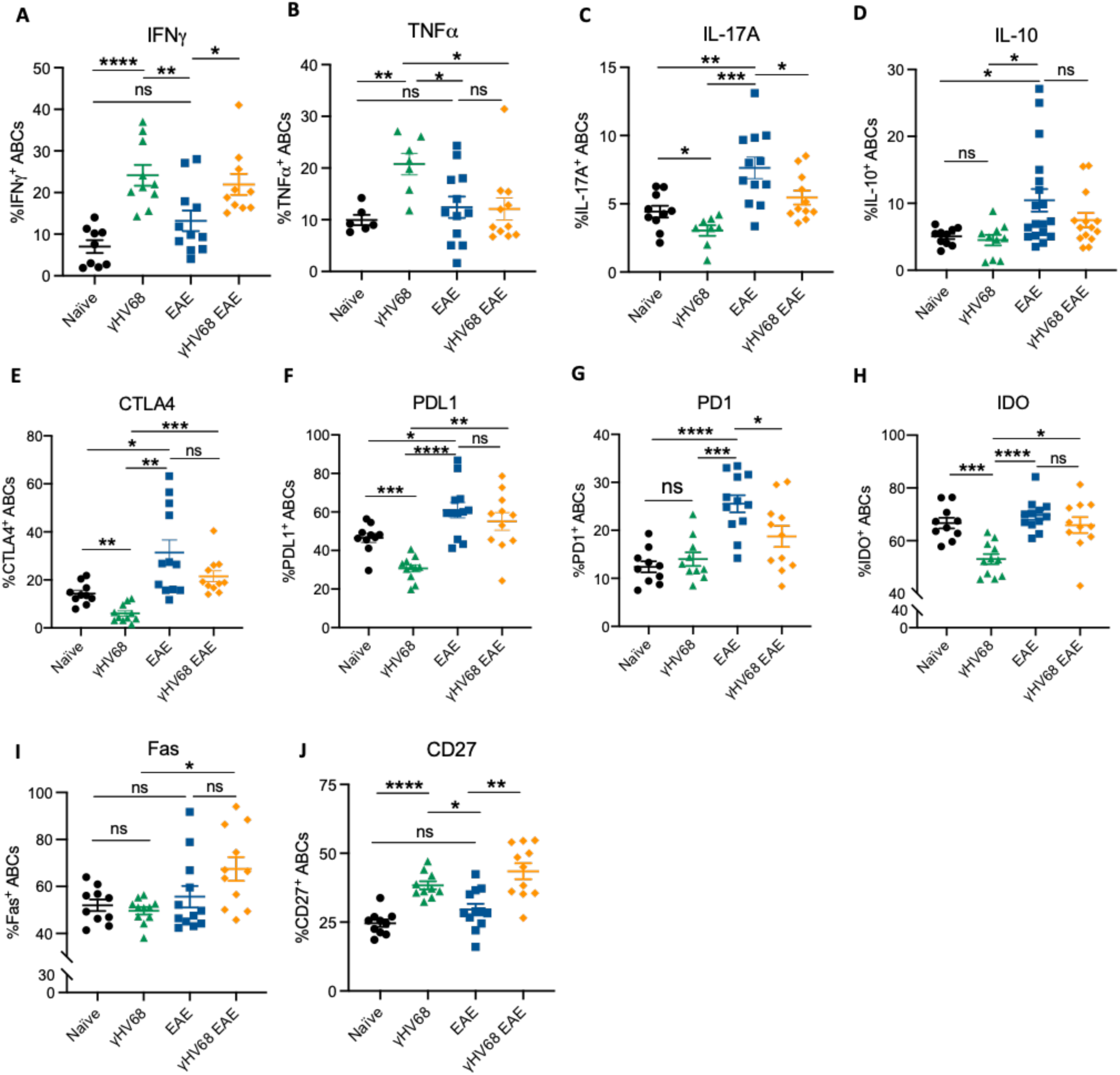
C57BL/6(J) mice infected with γHV68 for 35 days (γHV68, green triangles) or mock-infected (naïve, black circles). In some mice, MOG_35-55_ EAE was induced 35 days after mock infection (EAE, blue squares) or initial infection (γHV68 EAE, orange diamonds). At 35 days p.i. or 13 days post EAE induction, spleen was collected and processed for flow cytometry. Percentage of ABCs (CD19^+^CD11c^+^T-bet^+^) that are positive for (**A**) IFNγ, (**B**) TNF*α*, (**C**) IL-17A, (**D**) IL-10, (**E**) CTLA4, (**F**) PDL1, (**G**) PD1, (**H**) IDO, (**I**) Fas/CD95, and (**J**) CD27. Data in each graph are pooled across 2-3 experiments, n=9-18 per group. Data presented as mean ± SEM, analyzed by one-way ANOVA with multiple comparisons, ****p<0.0001, *** p<0.001, ** p<0.01, * p<0.05.

**Supplemental Figure 4:**
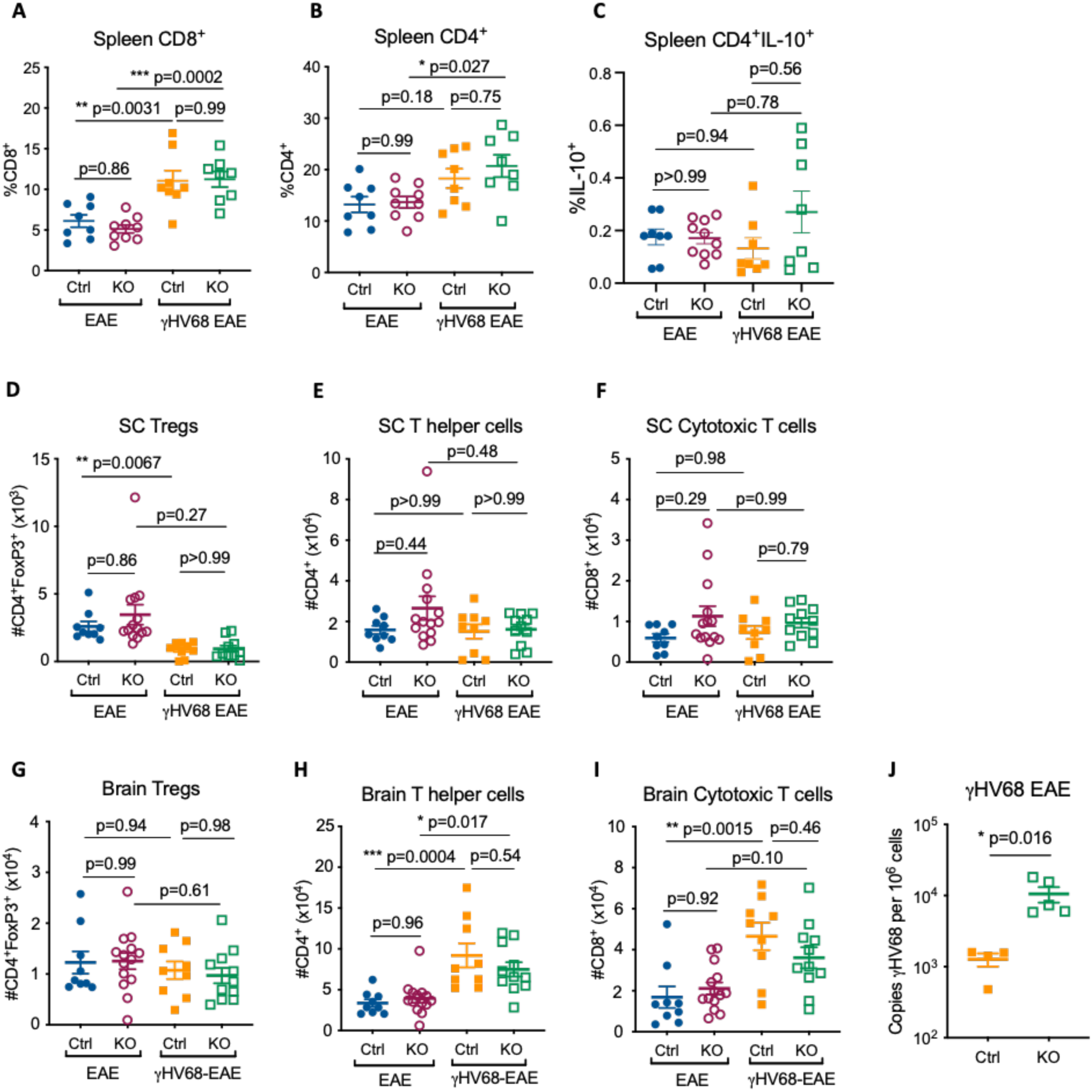
Female *Tbx21^fl/fl^Cd19^cre/+^ (*KO) and *Tbx21^fl/fl^Cd19^+/+^* (Ctrl) mice were infected with γHV68 or mock-infected for 35 days and then induced for MOG_35-55_ EAE. At day 13-18 post-EAE induction spleen, brain, and spinal cord collected. Spleen, brain, and spinal cord processed for flow cytometry and gDNA extracted from 4 × 10^6^ splenocytes. (**A**) Percent CD3^+^CD8^+^ (**A**) or CD3^+^CD4^+^ (**B**) of live lymphocytes in the spleen. (**C**) Percent of CD3^+^CD4^+^ T cells in the spleen that are IL-10^+^. (**D-F**) Total number, in the spinal cord, of CD45^+^CD3^+^CD4^+^FoxP3^+^ (**D**), CD45^+^CD3^+^CD4^+^ (**E**), and CD45^+^CD3^+^CD8^+^ (**F**) cells. (**G-I**) Total number, in the brain, of CD45^+^CD3^+^CD4^+^FoxP3^+^ (**G**), CD45^+^CD3^+^CD4^+^ (**H**), and CD45^+^CD3^+^CD8^+^ (**I**) cells. (**J**) Copies of γHV68 per 10^6^ splenocytes, as determined by qPCR. Analyzed by one-way ANOVA with multiple comparisons (**A-I**) or Mann-Whitney test (**J**), *** p<0.001, ** p<0.01, * p<0.05.

## References

1. Hao Y, O’Neill P, Naradikian MS, Scholz JL, Cancro MP. A B-cell subset uniquely responsive to innate stimuli accumulates in aged mice. Blood. 2011 Aug 4;118(5):1294–304.

2. Rubtsova K, Rubtsov AV, van Dyk LF, Kappler JW, Marrack P. T-box transcription factor T-bet, a key player in a unique type of B-cell activation essential for effective viral clearance. Proc Natl Acad Sci U S A. 2013 Aug 20;110(34):E3216–3224.

3. Rubtsov AV, Rubtsova K, Fischer A, Meehan RT, Gillis JZ, Kappler JW, et al. Toll-like receptor 7 (TLR7)-driven accumulation of a novel CD11c^+^ B-cell population is important for the development of autoimmunity. Blood. 2011 Aug 4;118(5):1305–15.

4. Ratliff M, Alter S, Frasca D, Blomberg BB, Riley RL. In senescence, age-associated B cells secrete TNFα and inhibit survival of B-cell precursors*. Aging Cell. 2013 Apr 1;12(2):303–11.

5. Rubtsov AV, Rubtsova K, Kappler JW, Jacobelli J, Friedman RS, Marrack P. CD11c-expressing B cells are located at the T cell B cell border in spleen and are potent antigen presenting cells. J Immunol Baltim Md 1950. 2015 Jul 1;195(1):71–9.

6. Claes N, Fraussen J, Vanheusden M, Hellings N, Stinissen P, Van Wijmeersch B, et al. Age-Associated B Cells with Proinflammatory Characteristics Are Expanded in a Proportion of Multiple Sclerosis Patients. J Immunol Baltim Md 1950. 2016 15;197(12):4576–83.

7. Isnardi I, Ng Y-S, Menard L, Meyers G, Saadoun D, Srdanovic I, et al. Complement receptor 2/CD21-human naive B cells contain mostly autoreactive unresponsive clones. Blood. 2010 Jun 17;115(24):5026–36.

8. Wehr C, Eibel H, Masilamani M, Illges H, Schlesier M, Peter H-H, et al. A new CD21low B cell population in the peripheral blood of patients with SLE. Clin Immunol Orlando Fla. 2004 Nov;113(2):161–71.

9. Rakhmanov M, Keller B, Gutenberger S, Foerster C, Hoenig M, Driessen G, et al. Circulating CD21low B cells in common variable immunodeficiency resemble tissue homing, innate-like B cells. Proc Natl Acad Sci U S A. 2009 Aug 11;106(32):13451–6.

10. Wang S, Wang J, Kumar V, Karnell JL, Naiman B, Gross PS, et al. IL-21 drives expansion and plasma cell differentiation of autoreactive CD11c hi T-bet + B cells in SLE. Nat Commun [Internet]. 2018 May 1 [cited 2018 Nov 2];9(1):1758. Available from: https://www.nature.com/articles/s41467-018-03750-7

11. Thorarinsdottir K, Camponeschi A, Jonsson C, Granhagen Önnheim K, Nilsson J, Forslind K, et al. CD21-/low B cells associate with joint damage in rheumatoid arthritis patients. Scand J Immunol. 2019 Aug;90(2):e12792.

12. Rubtsova K, Rubtsov AV, Thurman JM, Mennona JM, Kappler JW, Marrack P. B cells expressing the transcription factor T-bet drive lupus-like autoimmunity. J Clin Invest. 2017 May 12;127(4):1392–404.

13. Rubtsova K, Rubtsov AV, Cancro MP, Marrack P. Age-associated B cells: A T-bet dependent effector with roles in protective and pathogenic immunity. J Immunol Baltim Md 1950 [Internet]. 2015 Sep 1 [cited 2018 Nov 5];195(5):1933–7. Available from: https://www.ncbi.nlm.nih.gov/pmc/articles/PMC4548292/

14. Nath N, Prasad R, Giri S, Singh AK, Singh I. T-bet is essential for the progression of experimental autoimmune encephalomyelitis. Immunology. 2006 Jul;118(3):384–91.

15. Rubtsova K, Rubtsov AV, Halemano K, Li SX, Kappler JW, Santiago ML, et al. T Cell Production of IFNγ in Response to TLR7/IL-12 Stimulates Optimal B Cell Responses to Viruses. PloS One. 2016;11(11):e0166322.

16. Knox JJ, Buggert M, Kardava L, Seaton KE, Eller MA, Canaday DH, et al. T-bet+ B cells are induced by human viral infections and dominate the HIV gp140 response. JCI Insight. 2017 Apr 20;2(8).

17. Johnson JL, Rosenthal RL, Knox JJ, Myles A, Naradikian MS, Madej J, et al. The Transcription Factor T-bet Resolves Memory B Cell Subsets with Distinct Tissue Distributions and Antibody Specificities in Mice and Humans. Immunity [Internet]. 2020 Apr 29 [cited 2020 May 15]; Available from: http://www.sciencedirect.com/science/article/pii/S1074761320301333

18. Rubtsova K, Rubtsov AV, Dyk LF van, Kappler JW, Marrack P. T-box transcription factor T-bet, a key player in a unique type of B-cell activation essential for effective viral clearance. Proc Natl Acad Sci [Internet]. 2013 Aug 20 [cited 2018 Nov 2];110(34):E3216–24. Available from: http://www.pnas.org/content/110/34/E3216

19. A N, E Z, Cd S, Rg K, Cm T, Cf F, et al. Influenza-specific effector memory B cells predict long-lived antibody responses to vaccination in humans. 2019 May 20 [cited 2021 Jun 1]; Available from: https://europepmc.org/article/ppr/ppr79744

20. Austin JW, Buckner CM, Kardava L, Wang W, Zhang X, Melson VA, et al. Overexpression of T-bet in HIV infection is associated with accumulation of B cells outside germinal centers and poor affinity maturation. Sci Transl Med [Internet]. 2019 Nov 27 [cited 2021 Jun 1];11(520). Available from: https://stm.sciencemag.org/content/11/520/eaax0904

21. Chang L-Y, Li Y, Kaplan DE. Hepatitis C viraemia reversibly maintains subset of antigen-specific T-bet+ tissue-like memory B cells. J Viral Hepat [Internet]. 2017 May [cited 2020 May 15];24(5):389–96. Available from: https://www.ncbi.nlm.nih.gov/pmc/articles/PMC5637374/

22. Eccles JD, Turner RB, Kirk NA, Muehling LM, Borish L, Steinke JW, et al. T-bet+ Memory B Cells Link to Local Cross-Reactive IgG upon Human Rhinovirus Infection. Cell Rep [Internet]. 2020 Jan 14 [cited 2021 Jun 1];30(2):351–366.e7. Available from: https://www.sciencedirect.com/science/article/pii/S2211124719316778

23. Holla P, Dizon B, Ambegaonkar AA, Rogel N, Goldschmidt E, Boddapati AK, et al. Shared transcriptional profiles of atypical B cells suggest common drivers of expansion and function in malaria, HIV, and autoimmunity. Sci Adv [Internet]. 2021 May 26 [cited 2021 Jun 24];7(22):eabg8384. Available from: https://www.ncbi.nlm.nih.gov/pmc/articles/PMC8153733/

24. Cohen JI. Epstein–Barr Virus Infection. N Engl J Med. 2000 Aug 17;343(7):481–92.

25. Fraser KB, Millar JHD, Haire M, Mccrea S. INCREASED TENDENCY TO SPONTANEOUS IN-VITRO LYMPHOCYTE TRANSFORMATION IN CLINICALLY ACTIVE MULTIPLE SCLEROSIS. The Lancet. 1979 Oct 6;314(8145):715–7.

26. Ascherio A, Munger KL, Lennette ET, Spiegelman D, Hernán MA, Olek MJ, et al. Epstein-Barr virus antibodies and risk of multiple sclerosis: a prospective study. JAMA. 2001 Dec 26;286(24):3083–8.

27. Farrell RA, Antony D, Wall GR, Clark DA, Fisniku L, Swanton J, et al. Humoral immune response to EBV in multiple sclerosis is associated with disease activity on MRI. Neurology [Internet]. 2009 Jul 7 [cited 2019 Mar 22];73(1):32–8. Available from: https://www.ncbi.nlm.nih.gov/pmc/articles/PMC2848585/

28. Pender MP, Csurhes PA, Burrows JM, Burrows SR. Defective T-cell control of Epstein–Barr virus infection in multiple sclerosis. Clin Transl Immunol [Internet]. 2017 Jan 20 [cited 2019 Mar 22];6(1):e126. Available from: https://www.ncbi.nlm.nih.gov/pmc/articles/PMC5292561/

29. James JA, Kaufman KM, Farris AD, Taylor-Albert E, Lehman TJ, Harley JB. An increased prevalence of Epstein-Barr virus infection in young patients suggests a possible etiology for systemic lupus erythematosus. J Clin Invest [Internet]. 1997 Dec 15 [cited 2021 Jun 1];100(12):3019–26. Available from: https://www.ncbi.nlm.nih.gov/pmc/articles/PMC508514/

30. James JA, Neas BR, Moser KL, Hall T, Bruner GR, Sestak AL, et al. Systemic lupus erythematosus in adults is associated with previous Epstein-Barr virus exposure. Arthritis Rheum. 2001 May;44(5):1122–6.

31. Li Z-X, Zeng S, Wu H-X, Zhou Y. The risk of systemic lupus erythematosus associated with Epstein-Barr virus infection: a systematic review and meta-analysis. Clin Exp Med. 2019 Feb;19(1):23–36.

32. Balandraud N, Meynard JB, Auger I, Sovran H, Mugnier B, Reviron D, et al. Epstein-Barr virus load in the peripheral blood of patients with rheumatoid arthritis: accurate quantification using real-time polymerase chain reaction. Arthritis Rheum. 2003 May;48(5):1223–8.

33. Ferrell PB, Aitcheson CT, Pearson GR, Tan EM. Seroepidemiological study of relationships between Epstein-Barr virus and rheumatoid arthritis. J Clin Invest. 1981 Mar;67(3):681–7.

34. Alspaugh MA, Henle G, Lennette ET, Henle W. Elevated levels of antibodies to Epstein-Barr virus antigens in sera and synovial fluids of patients with rheumatoid arthritis. J Clin Invest. 1981 Apr;67(4):1134–40.

35. Márquez AC, Horwitz MS. The Role of Latently Infected B Cells in CNS Autoimmunity. Front Immunol [Internet]. 2015 Oct 28 [cited 2017 Apr 5];6. Available from: http://www.ncbi.nlm.nih.gov/pmc/articles/PMC4623415/

36. Bar-Or A, Pender MP, Khanna R, Steinman L, Hartung H-P, Maniar T, et al. Epstein-Barr Virus in Multiple Sclerosis: Theory and Emerging Immunotherapies. Trends Mol Med. 2020 Mar;26(3):296–310.

37. Olivadoti M, Toth LA, Weinberg J, Opp MR. Murine gammaherpesvirus 68: a model for the study of Epstein-Barr virus infections and related diseases. Comp Med. 2007 Feb;57(1):44–50.

38. Casiraghi C, Shanina I, Cho S, Freeman ML, Blackman MA, Horwitz MS. Gammaherpesvirus Latency Accentuates EAE Pathogenesis: Relevance to Epstein-Barr Virus and Multiple Sclerosis. PLOS Pathog [Internet]. 2012 May 17 [cited 2018 Dec 13];8(5):e1002715. Available from: https://journals.plos.org/plospathogens/article?id=10.1371/journal.ppat.1002715

39. Casiraghi C, Márquez AC, Shanina I, Horwitz MS. Latent virus infection upregulates CD40 expression facilitating enhanced autoimmunity in a model of multiple sclerosis. Sci Rep. 2015 Sep 10;5:srep13995.

40. Lau D, Lan LY-L, Andrews SF, Henry C, Rojas KT, Neu KE, et al. Low CD21 expression defines a population of recent germinal center graduates primed for plasma cell differentiation. Sci Immunol. 2017 Jan 27;2(7).

41. Barnett BE, Staupe RP, Odorizzi PM, Palko O, Tomov VT, Mahan AE, et al. B cell intrinsic T-bet expression is required to control chronic viral infection. J Immunol Baltim Md 1950 [Internet]. 2016 Aug 15 [cited 2018 Nov 13];197(4):1017–22. Available from: https://www.ncbi.nlm.nih.gov/pmc/articles/PMC4975981/

42. Rubtsova K, Marrack P, Rubtsov AV. Age-associated B cells: are they the key to understanding why autoimmune diseases are more prevalent in women? Expert Rev Clin Immunol [Internet]. 2012 Jan [cited 2021 Jun 1];8(1):5–7. Available from: https://www.ncbi.nlm.nih.gov/pmc/articles/PMC3940262/

43. Palaszynski KM, Loo KK, Ashouri JF, Liu H, Voskuhl RR. Androgens are protective in experimental autoimmune encephalomyelitis: implications for multiple sclerosis. J Neuroimmunol. 2004 Jan;146(1–2):144–52.

44. Okuda Y, Okuda M, Bernard CCA. Gender does not influence the susceptibility of C57BL/6 mice to develop chronic experimental autoimmune encephalomyelitis induced by myelin oligodendrocyte glycoprotein. Immunol Lett. 2002 Apr 1;81(1):25–9.

45. Mouat IC, Morse ZJ, Shanina I, Brown KL, Horwitz MS. Latent gammaherpesvirus exacerbates arthritis through modification of age-associated B cells. Valenzano DR, editor. eLife [Internet]. 2021 Jun 3 [cited 2021 Jun 6];10:e67024. Available from: https://doi.org/10.7554/eLife.67024

46. Molnarfi N, Schulze-Topphoff U, Weber MS, Patarroyo JC, Prod’homme T, Varrin-Doyer M, et al. MHC class II-dependent B cell APC function is required for induction of CNS autoimmunity independent of myelin-specific antibodies. J Exp Med. 2013 Dec 16;210(13):2921–37.

47. Goncalves DaSilva A, Liaw L, Yong VW. Cleavage of Osteopontin by Matrix Metalloproteinase-12 Modulates Experimental Autoimmune Encephalomyelitis Disease in C57BL/6 Mice. Am J Pathol [Internet]. 2010 Sep [cited 2021 Jun 2];177(3):1448–58. Available from: https://www.ncbi.nlm.nih.gov/pmc/articles/PMC2928976/

48. Doss PMIA, Umair M, Baillargeon J, Fazazi R, Fudge N, Akbar I, et al. Male sex chromosomal complement exacerbates the pathogenicity of Th17 cells in a chronic model of central nervous system autoimmunity. Cell Rep. 2021 Mar 9;34(10):108833.

49. Márquez AC, Shanina I, Horwitz MS. Multiple Sclerosis-Like Symptoms in Mice Are Driven by Latent γHerpesvirus-68 Infected B Cells. Front Immunol [Internet]. 2020 [cited 2021 Apr 12];11. Available from: https://www.frontiersin.org/articles/10.3389/fimmu.2020.584297/full

